# Depression of substantia nigra dopamine transmission is driven by retrograde neurotensin release and is enhanced by methamphetamine self-administration

**DOI:** 10.1101/717843

**Authors:** Christopher W. Tschumi, Ramaswamy Sharma, William B. Lynch, Amanda L. Sharpe, Michael J. Beckstead

## Abstract

Midbrain dopamine neurons play central roles in reward learning and motivated behavior, and inhibition at somatodendritic dopamine D2 receptor (D2R) synapses blunts psychostimulant reinforcement. Release of the neuropeptide neurotensin in the midbrain increases following methamphetamine exposure and induces long-term depression of D2R synaptic currents (LTD_DA_), however the source of neurotensin that drives LTD_DA_ is not known. Here we show that LTD_DA_ is driven by neurotensin released by dopamine neurons. Optogenetic stimulation of dopamine neurons was sufficient to induce LTD_DA_ in the substantia nigra, but not the ventral tegmental area, and was dependent on neurotensin receptors, postsynaptic calcium, and vacuolar-type H^+^-ATPase activity in the postsynaptic cell. Further, LTD_DA_ was enhanced in mice that had self-administered methamphetamine. These findings reveal a novel form of signaling between dopamine neurons involving release of the peptide neurotensin, which may act as a feed forward mechanism to increase dopamine neuron excitability and methamphetamine self-administration.

## Introduction

Midbrain dopamine neurons of the substantia nigra pars compacta (SNc) and ventral tegmental area (VTA) play central roles in the initiation of movement and motivated behavior. Thus, a complete understanding of the determinants of dopamine neuron excitability is of critical importance in understanding these complex behaviors. Dopamine neurons inhibit each other through a relatively slow form of dendritic neurotransmission mediated by postsynaptic dopamine D2 autoreceptors (D2Rs, Beckstead et al; 2004; Robinson et al., 2017). Dendritic dopamine neurotransmission can be measured in mouse brain slices by electrically evoking dopamine release and electrophysiologically recording the subsequent D2R-, potassium channel-mediated inhibitory postsynaptic currents (D2-IPSCs, Beckstead et al., 2004). Previous work from our lab shows that D2-IPSCs undergo long-term depression (LTD_DA_) subsequent to low-frequency electrical stimulation (Beckstead and Williams, 2007; Piccart et al., 2015) through a mechanism likely involving signaling by the neuropeptide neurotensin (NT, Piccart et al., 2015).

NT is expressed throughout the central nervous system, and in both the midbrain and striatum (a major dopamine projection region) it can affect dopamine-mediated behaviors including food- and psychostimulant-reinforced responding (Dominguez-Lopez et al., 2018; Kempadoo et al., 2013; Leinninger et al., 2011; Patterson et al., 2015; Tschumi and Beckstead, 2018; Tschumi and Beckstead, 2019). Conversely, exposure to abused drugs including morphine, cocaine, and methamphetamine can induce NT peptide upregulation or release across different brain regions (Betancur et al., 2001; Frankel et al., 2011; Geisler and Zahm, 2006; Hanson et al., 2013; Stiller et al., 1997; Tschumi and Beckstead, 2019). Incredibly, exogenous NT application induces synaptic plasticity of GABAergic, dopaminergic, and glutamatergic input to dopamine neurons, as well as activating cell-autonomous endocannabinoid signaling in those cells (Kortleven et al., 2012; Kempadoo et al., 2013; Bose et al., 2015; Stuhrman and Roseberry, 2015; Piccart et al., 2015; Gantz and Bean, 2017; Tschumi and Beckstead, 2018; Tschumi and Beckstead, 2019). NT release from lateral hypothalamic inputs to the VTA drives plasticity at glutamatergic synapses (Kempadoo et al., 2013), but the endogenous source of NT which drives plasticity at dendrodendritic dopamine synapses is not known. Therefore, physiological delineation of NT signaling is central to understanding synaptic plasticity in, and excitability of, dopamine neurons.

Here we show that LTD_DA_ in the SNc is driven by neurotensin release from dopamine neurons. We used transgenic and viral targeting to enable optogenetic stimulation of dopamine neurons or NT terminals projecting from either the lateral hypothalamus (LH) or nucleus accumbens (NAc) to the VTA. Targeted stimulation of dopamine neurons drove LTD_DA_ in the SNc that was prevented by neurotensin receptor antagonism, as well as postsynaptic chelation of calcium or inhibition of vacuolar-type H^+^-ATPase (V-ATPase). LTD_DA_ was also enhanced in mice that self-administer methamphetamine, however, neither stimulation of dopaminergic, lateral hypothalamic, or accumbal NT-expressing inputs were sufficient to drive LTD_DA_ in the VTA. These findings provide evidence for a second form of inter-dopamine neuron signaling which proceeds through local NT release, and indicates that this form of communication may be dysregulated as a result of methamphetamine use.

## Results

### Repetitive stimulation of dopamine neurons is sufficient to drive LTD_DA_ in the SNc

We previously found that stimulation-induced long-term depression of D2-IPSCs (LTD_DA_, Beckstead and Williams, 2007) is driven by endogenous NT signaling (Piccart et al., 2015) and suggested that dopamine neurons might also release NT. Thus, to determine the source of NT which drives LTD_DA_ we used a combination of transgenic and optogenetic tools to delineate the role of dopamine neurons and NT afferents to the midbrain. We first generated a mouse expressing channelrhodopsin (ChR2) specifically in dopamine neurons by cross-breeding dopamine transporter (DAT)-Cre and cre-dependent ChR2 mouse lines (Figure 1 A1-3, B). Using whole-cell voltage clamp (−55 mV) recordings of SNc dopamine neurons, we determined that blue light optical stimulation, like electrical stimulation (Beckstead et al., 2004), generated slow outward synaptic currents that were blocked by the dopamine D2 receptor antagonist sulpiride (D2-oIPSCs, Figure 1 C1-2) or by the voltage-gated sodium channel blocker tetrodotoxin (Figure 1 D), and were dependent on the presence of extracellular Ca^2+^ (Figure 1 E). This suggests that optogenetic excitation of dopamine neurons and subsequent dopamine release closely resembles physiological dopamine neurotransmission (Beckstead et al., 2004). Furthermore, acute bath application of methamphetamine increased both D2-oIPSC amplitude and width, suggesting that the kinetics of the synaptic current are tightly regulated by dopamine uptake (Figure 1 F1-3). Thus, optogenetic stimulation allows for an assessment of dopamine neuron excitation without stimulation of other cell types and without including blockers of other neurotransmitter receptors, and revealed that repeated optostimulation of dopamine neurons (10 Hz, 2 min) is sufficient to drive LTD_DA_ (t_5_ = 5.04, p = 0.0040; Figure 1 G1-3).

**Figure 1.**
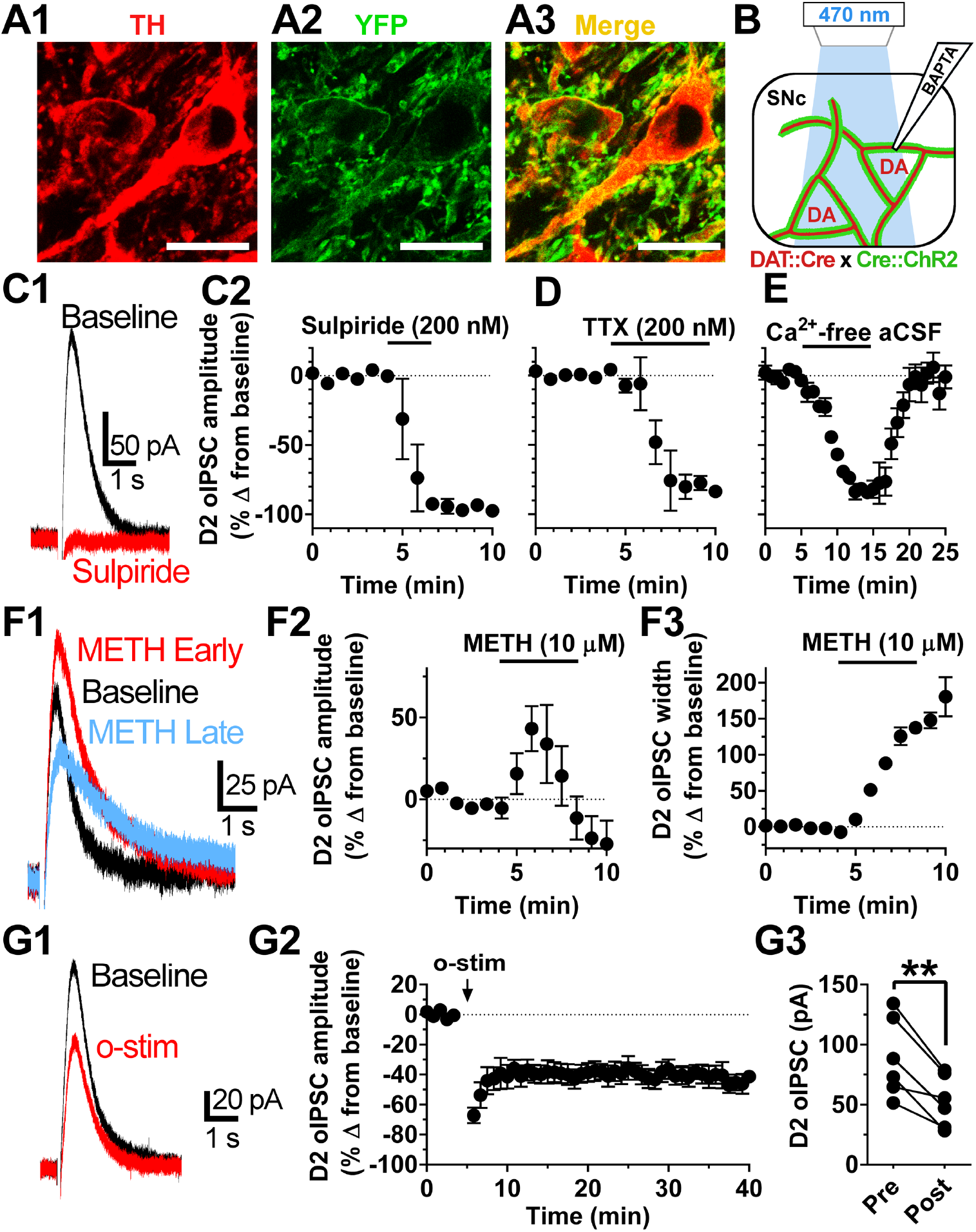
Stimulation of dopamine neurons is sufficient to evoke D2-IPSCs and drive LTD_DA_ in the SNc. Images depicting tyrosine hydroxylase (TH) (A1), eYFP-fused to ChR2 (A2), and co-localization (A3), scale bar = 20 µm. Schematic depicting dopamine (DA) neurons from DAT-Cre mice (DAT::Cre) crossed with Ai32 (Cre::ChR2) mice resulting in dopamine neuron-specific expression of eYFP-fused to ChR2 and stimulation by blue light in the SNc (470 nm, B). A blue-light elicited D2-oIPSC during baseline and following bath perfusion of sulpiride (C1), and time-course (C2). Time-courses depicting reduction of D2-oIPSC amplitude during bath perfusion of TTX (D) and calcium-free aCSF (E). Effects of methamphetamine (METH) on D2-oIPSCs (during the peak-effect of METH on amplitude, and later when the effect of METH on half-width was more pronounced, F1), time-course of effect on amplitude (F2), and half-width (F3). Effect of dopamine-neuron specific stimulation (o-stim, 10 Hz blue light for 2 min) on D2-oIPSCs (G1), time-course (G2), and raw amplitudes (G3) during an average of 5 baseline sweeps (Pre) and the 5 sweeps 19-23 minutes following optostimulation (Post). **p < 0.01

### LTD_DA_ requires NT type-2 receptor signaling, and postsynaptic calcium and V-ATPase

As dopamine neurons express both type 1 and 2 NT receptors (Sarret et al., 1998; Mazella et al., 1988), we next investigated if optostimulation-induced LTD_DA_ is dependent on NT signaling. Pre-incubation with the non-selective NT type 1/2 receptor antagonist SR 142948 prevented LTD_DA_ (Figure 2 A 1-2, F; one-way ANOVA, F_4,26_ = 18.92, p < 0.0001, Tukey’s, p = 0.0001). Conversely and consistent with our previous findings (Piccart et al., 2015), a selective NT type-1 receptor antagonist failed to prevent LTD_DA_ (Figure 2 B 1-2, F; p = 0.99). As presumably only dopamine neurons are being depolarized by the blue light, this suggests that dopamine neurons are themselves releasing NT that induces LTD_DA_ though activation of NT type-2 receptors. Furthermore, buffering calcium by dialyzing the postsynaptic cell with BAPTA (10 mM) added to the recording pipette prevented optostimulation-induced LTD_DA_ (Figure 2 C 1-2, F; p = 0.027), but not depression of D2-oIPSCs induced by bath application of the active fragment of NT (NT_8-13_, Figure 2 D 1-2, t_3_ = 4.1, p = 0.026). Together, these findings suggest that NT-induced depression of D2-oIPSCs is not dependent on intracellular calcium *per se*, as NT perfusion may bypass a postsynaptic calcium-dependent step in stimulation-induced LTD_DA_.

**Figure 2.**
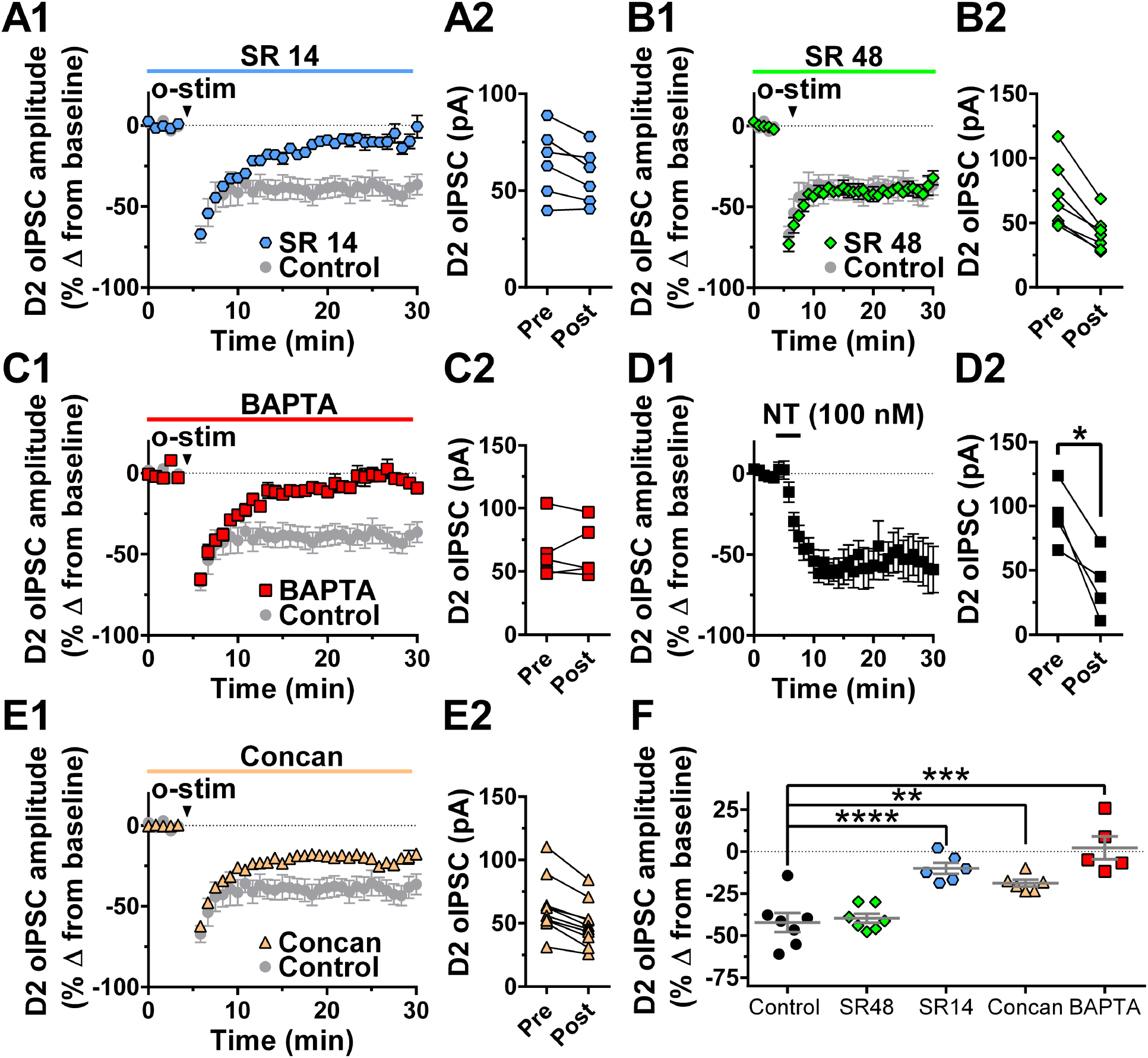
LTD_DA_ requires NT type-2 receptor signaling, plus postsynaptic calcium and V-ATPase. Effect of dopamine neuron-specific stimulation (o-stim, 10 Hz blue light for 2 min) on D2-oIPSCs in aCSF (control, grey circles, data first shown in Figure 1) or in the presence of: the non-selective NT type-1/2 receptor antagonist SR142948 (1 µM, SR 14, blue hexagons), time-course (A1) and raw amplitudes (A2); the NT type-1 receptor antagonist SR48692 (green diamonds, B1, B2); or with BAPTA (10 mM) added to the recording pipette internal solution (red squares, C1, C2). The effect of 5-minute bath perfusion of NT_8-13_ on D2-oIPSCs with BAPTA (10 mM) in the recording pipette (D1, D2). Effect of optostimulation on D2-oIPSCs with concanamycin A (5 µM) in the recording pipette (E1, E2). Comparison of percent change from baseline amplitude of D2-oIPSCs following optostimulation between groups (F). All raw amplitudes and percent changes are the average of 5 baseline sweeps (Pre) and the 5 sweeps 19-23 minutes following optostimulation (Post). Arrowhead notates optostimulation of dopamine neurons (10 Hz, 2 min). *p < 0.05, **p < 0.01, ***p < 0.001

NT-induced synaptic plasticity in other brain regions is dependent on dopamine D1 receptor (D1R) signaling (Krawczyk et al., 2013), and an alternative explanation is that D1R-expressing axon terminals in the midbrain (Levey et al., 1993) could produce NT release subsequent to optostimulation of dopamine neurons. However, the selective D1R antagonist SKF 38393 (10 µM) failed to prevent LTD_DA_ (Figure S1 A, B; t_4_ = 5.5, p = 0.0054). As calcium chelation prevents optostimulation-induced LTD_DA_, and NT is released from dopamine neurons in a calcium-dependent manner (Kitabgi et al., 1990), we next investigated if the dopamine neuron being recorded releases NT to drive LTD_DA_ in a cell-autonomous manner. NT is found in dense-core vesicles in dopamine neuron dendrites (Bayer et al., 1991), release of which should be inhibited by V-ATPase, which is necessary for both proper trafficking of neuropeptide-containing dense core vesicles (Taupenot et al., 2005) and NT release in other brain regions (Krawczyk et al., 2013, Normandeau et al., 2017). Indeed, dialyzing the postsynaptic cell with the V-ATPase inhibitor concanamycin A partially blunted LTD_DA_ (Figure 2 E 1-2, F; p = 0.005). While these experiments do not definitively exclude the release of other signaling molecules, these results suggest that NT is released in a calcium- and V-ATPase-dependent manner from dopamine neurons located postsynaptic to other dopamine neurons.

### Repetitive stimulation of dopamine neurons or other NT inputs is not sufficient to drive LTD_DA_ in the VTA

As NT peptide has been observed in cell bodies and terminals in both the SNc and VTA, we next examined if dopamine neuron stimulation is sufficient to drive LTD_DA_ in VTA dopamine neurons. Although we have previously shown that bath perfusion of NT induces depression of D2R signaling in VTA dopamine neurons (Piccart et al., 2015), optostimulation surprisingly failed to induce LTD_DA_ in most of the VTA neurons recorded (Figure 3 A 1-3; t_4_ = 0.75, p = 0.50), suggesting the presence of brain region-specific interactions between NT and dopamine transmission. However, consistent with previous findings (Beckstead and Williams, 2007), electrical stimulation did induce LTD_DA_ in the VTA (Figure S2 A, B). The fact that optostimulation-induced LTD_DA_ is NT-dependent combined with previous findings that bath-perfused NT depresses D2-IPSCs (Piccart et al., 2015) suggests that electrical stimulation may elicit NT release from non-dopamine cells to induce LTD_DA_ in VTA dopamine neurons.

**Figure 3.**
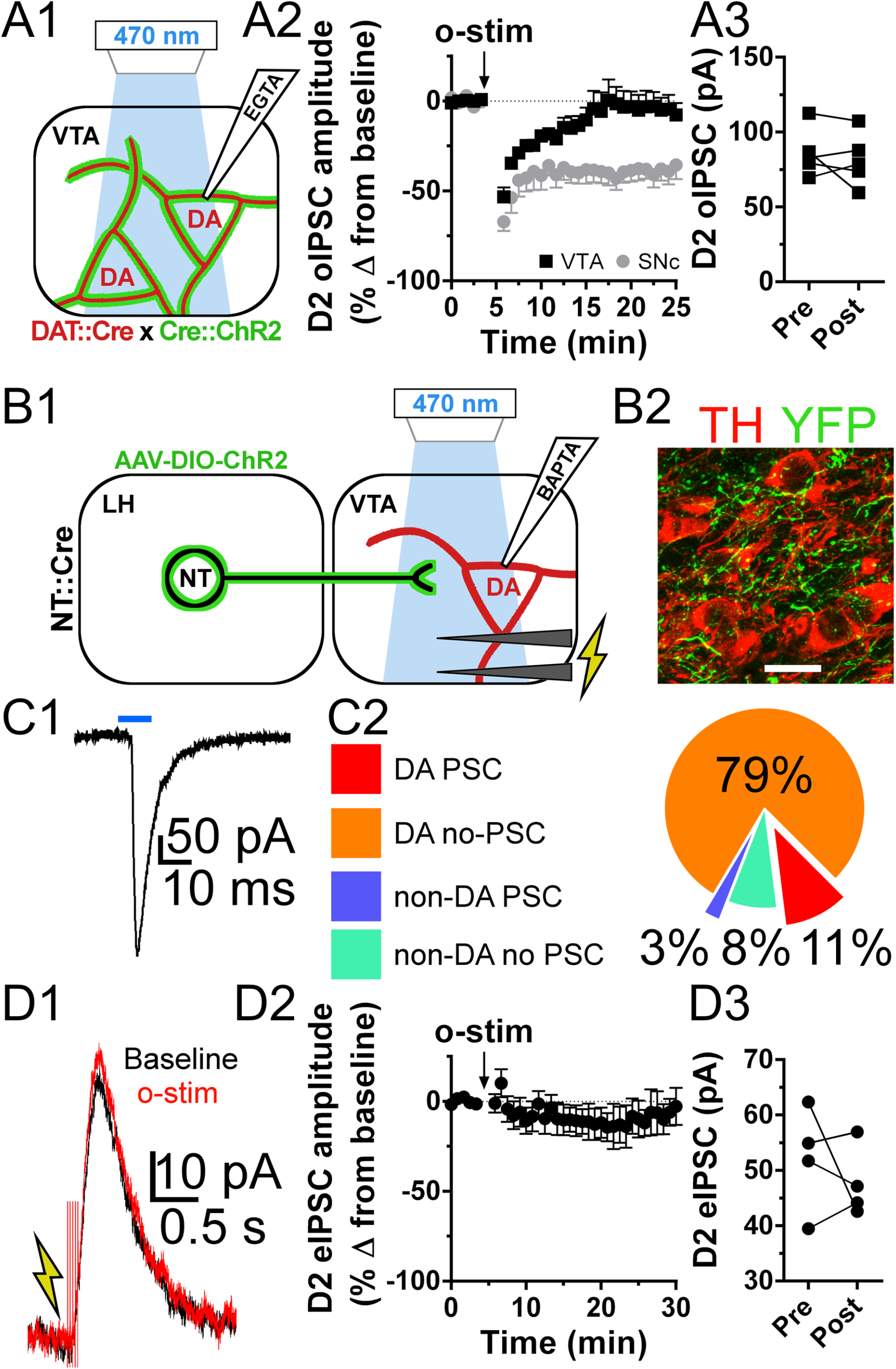
Stimulation of dopamine neurons or NT-expressing inputs from the LH to the VTA does not drive LTD_DA_. Schematic depicting dopamine (DA) neurons from DAT-Cre mice (DAT::Cre) crossed with Ai32 (Cre::ChR2) mice resulting in dopamine neuron-specific expression of eYFP-fused to ChR2 and stimulation by blue light in the VTA (470 nm, A1). Effect of dopamine-neuron specific stimulation (o-stim, 10 Hz blue light for 2 min) on D2-oIPSCs recorded from SNc (SNc, grey circles, data first shown in Figure 1) or VTA dopamine neurons (VTA, black squares), time-course (A2) and raw amplitudes (A3). Schematic depicting NT-expressing neurons in NTS-Cre mice following intra-LH injection of an AAV that drives cre-inducible expression of eYFP-fused to ChR2 (AAV-DIO-ChR2). These inputs were stimulated either by single pulses of blue light (10 ms, 470 nm) or repeated blue-light stimulation (10 Hz, 2 min, 5 ms pulses) to drive NT release (B1). Electrical stimulation (with GABA and glutamate antagonists in the aCSF) was used to elicit D2-eIPSCs. Sample image depicting expression of eYFP-fused ChR2 (YFP) near tyrosine-hydroxylase (TH) containing neurons (B2, scale bar = 20 µm). Sample trace of a postsynaptic current (PSC) generated by LH-NT input stimulation (C1, blue bar = 10 ms blue-light pulse). Chart depicting the percent of dopamine (DA) and non-dopamine neurons (non-DA) in which a PSC was or was not (no PSC) elicited by single-pulse stimulation (N = 38, C2). Effect of repeated stimulation of LH NT input to VTA (o-stim, 10 Hz blue light for 2 min) on D2-eIPSCs, sample trace (D1), time-course (D2), and raw amplitudes (D3). Raw amplitudes are the average 5 baseline sweeps (Pre) and 5 sweeps 19-23 minutes following optostimulation (Post).

We next investigated two prominent NT-expressing inputs to the VTA: 1) the LH afferents to the VTA, which have only weak projection to the SNc (Patterson et al., 2015), and 2) the NAc inputs to the VTA, which express increased levels of NT following methamphetamine exposure (Geisler and Zahm, 2006). NT, like many neuropeptides, is co-expressed in neurons that form synaptic connections and generate either GABA- or glutamate-mediated postsynaptic currents in VTA dopamine neurons (Leinninger et al., 2013; Kempadoo et al., 2013; Tschumi and Beckstead, 2019). In order to specifically stimulate these inputs, NTS-Cre mice that express Cre-recombinase specifically in cells that express NT were administered bilateral intra-LH or intra-NAc injections of AAV to drive cre-inducible expression of ChR2 (schematic; Figure 3 B1, 4 A1). Both immunohistochemistry (Figure 3 B2) and live-slice imaging (observed during recordings, data not shown) revealed terminals from NT-expressing LH inputs to the VTA, and optostimulation of these inputs generated fast postsynaptic currents in a fraction of both dopamine and non-dopamine neurons recorded (Figure 3 C1-2). However, recordings of D2-IPSCs in a separate group of cells revealed no induction of LTD_DA_ by targeted stimulation of NT-expressing LH inputs to the VTA (Figure 3 D1-3; t_3_ = 0.80, p = 0.48). Similarly, optogenetic stimulation of NT-expressing NAc inputs to the VTA (Figure 4 A1-2) generated postsynaptic currents in only a fraction of cells recorded (Figure 4 B1-2) and failed to induce LTD_DA_ (Figure 4 C1-3; t_7_ = 0.223, p = 0.87). Together, these results suggest that unlike in the SNc, LTD_DA_ in the VTA is not driven by NT release by dopamine neurons, nor by two major NT inputs arising from the NAc and LH.

**Figure 4.**
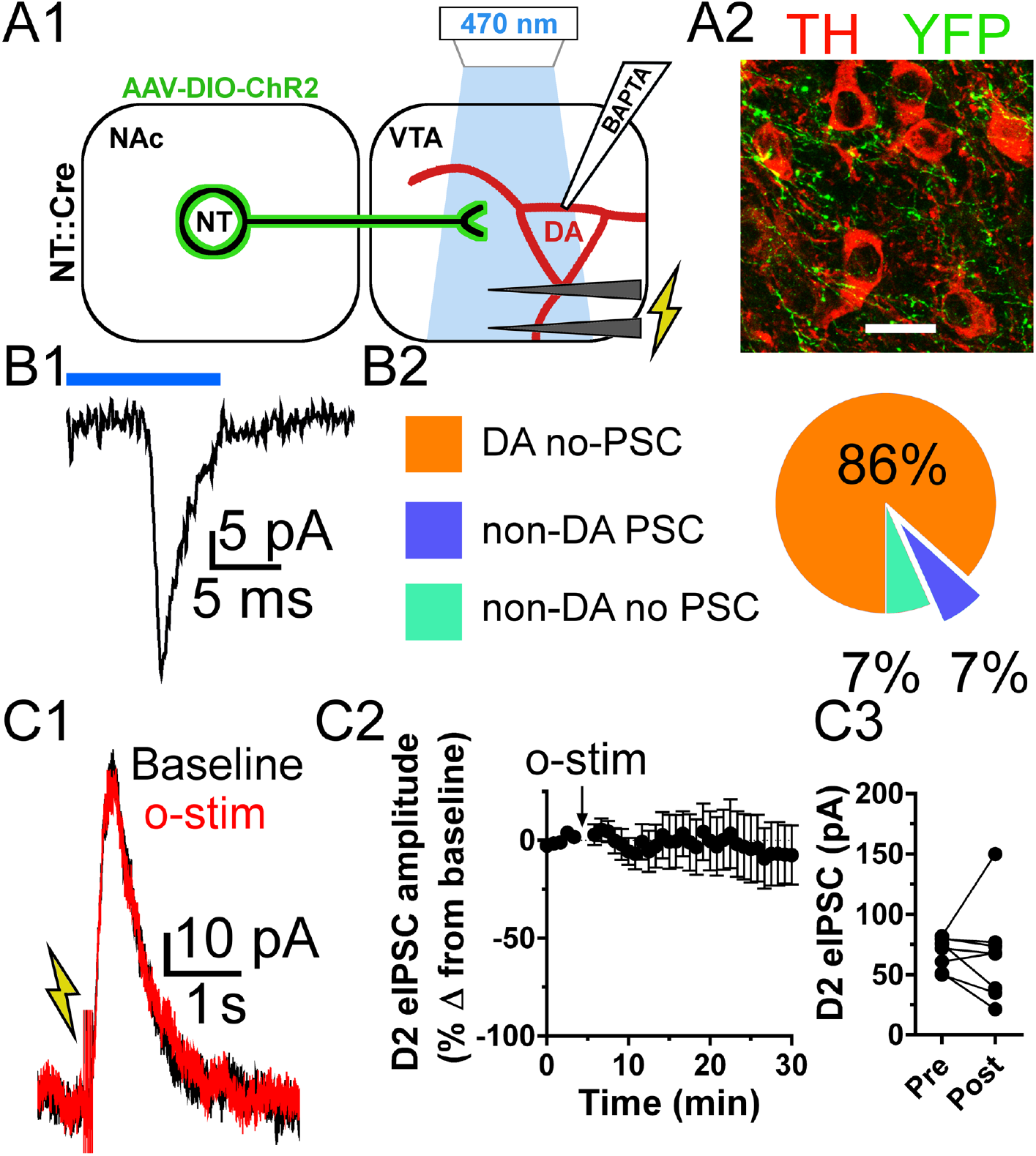
NT-expressing inputs from the NAc to the VTA do not drive LTD_DA_. Schematic depicting NT-expressing neurons in NTS-Cre mice following intra-NAc infusion of an AAV that drives cre-inducible expression of eYFP-fused to ChR2 (AAV-DIO-ChR2). These inputs were stimulated either by single pulses of blue light (10 ms, 470 nm) or repeated blue-light stimulation (10 Hz, 2 min, 5 ms pulses) to drive NT release (A1). Electrical stimulation (with GABA and glutamate antagonists in aCSF) was used to elicit D2-eIPSCs. Sample image depicting expression of eYFP-fused ChR2 (YFP) near tyrosine-hydroxylase (TH) containing neurons (A2, scale bar = 20 µm). Sample trace of a postsynaptic current (PSC) generated by LH-NT input stimulation (B1, blue bar = 10 ms blue-light pulse). Chart depicting the percent of dopamine (DA) and non-dopamine neurons (non-DA) in which a PSC was or was not (no PSC) elicited by single-pulse stimulation (N = 15, B2). Effect of repeated stimulation of NAc NT input to VTA (o-stim, 10 Hz blue light for 2 min) on D2-eIPSCs, sample trace (C1), time-course (C2), and raw amplitudes (C3). Raw amplitudes are the average 5 baseline sweeps (Pre) and 5 sweeps 19-23 minutes following optostimulation (Post).

### LTD_DA_ is enhanced in mice that self-administer methamphetamine

Midbrain dopamine signaling is perturbed presynaptically at the level of spontaneous dopamine release and postsynaptically at D2Rs following in vivo psychostimulant exposure (Gantz et al., 2015, Sharpe et al., 2014). To determine if dendrodendritic dopamine neurotransmission and plasticity are affected by repeated psychostimulant exposure, we measured D2-eIPSCs from mice that had a history of operant self-administration of methamphetamine. Following recovery from jugular catheter implantation surgery, mice were placed in an operant chamber for self-administration of methamphetamine (2 hours/day). Each operant chamber was equipped with 2 nose-poke holes, and responding in one hole resulted in an infusion of methamphetamine (0.1 mg/kg) while responding in the inactive hole had no consequence. Initially, mice were trained to self-administer methamphetamine on a schedule of reinforcement that progressed within-session from fixed ratio 1 (each response results in an infusion, FR1) to fixed ratio 3 (FR3). After mice acquired stable responding with high accuracy in the active hole (89.5 ± 4.0 %) they were transitioned to an FR3 schedule of reinforcement for a minimum of nine days. Mice self-administered 2.38 ± 0.10 mg/kg/day and continued to respond with a high degree of accuracy (91 ± 1.0% nose-pokes in active nose-poke hole) in the 5 sessions preceding electrophysiological recordings (Figure 5 A1-2). In recordings from SNc dopamine neurons in the presence of a cocktail of antagonists to block acetylcholine, GABA, and glutamate signaling, repetitive electrical stimulation (10 Hz, 2 min) induced LTD_DA_ in both groups (Figure 5 B1, B3; naïve, t_4_ = 4.86, p = 0.008; methamphetamine, t_6_ = 4.58, p = 0.004), however the effect was larger in neurons from mice that self-administered methamphetamine (Figure 5 B2; t_10_ = 2.34, p = 0.041). In agreement with previous studies from our lab showing that methamphetamine self-administration decreases D2R currents (Sharpe et al., 2014), D2-eIPSCs were of smaller amplitude in mice that self-administered methamphetamine compared to drug-naïve controls (Figure 5 B3; Naïve vs. METH; t_10_ = 3.10, p = 0.011). Together, these results suggest that although methamphetamine-self administration decreases dendritic dopamine transmission, the ability to induce LTD_DA_ persists and is actually enhanced.

**Figure 5.**
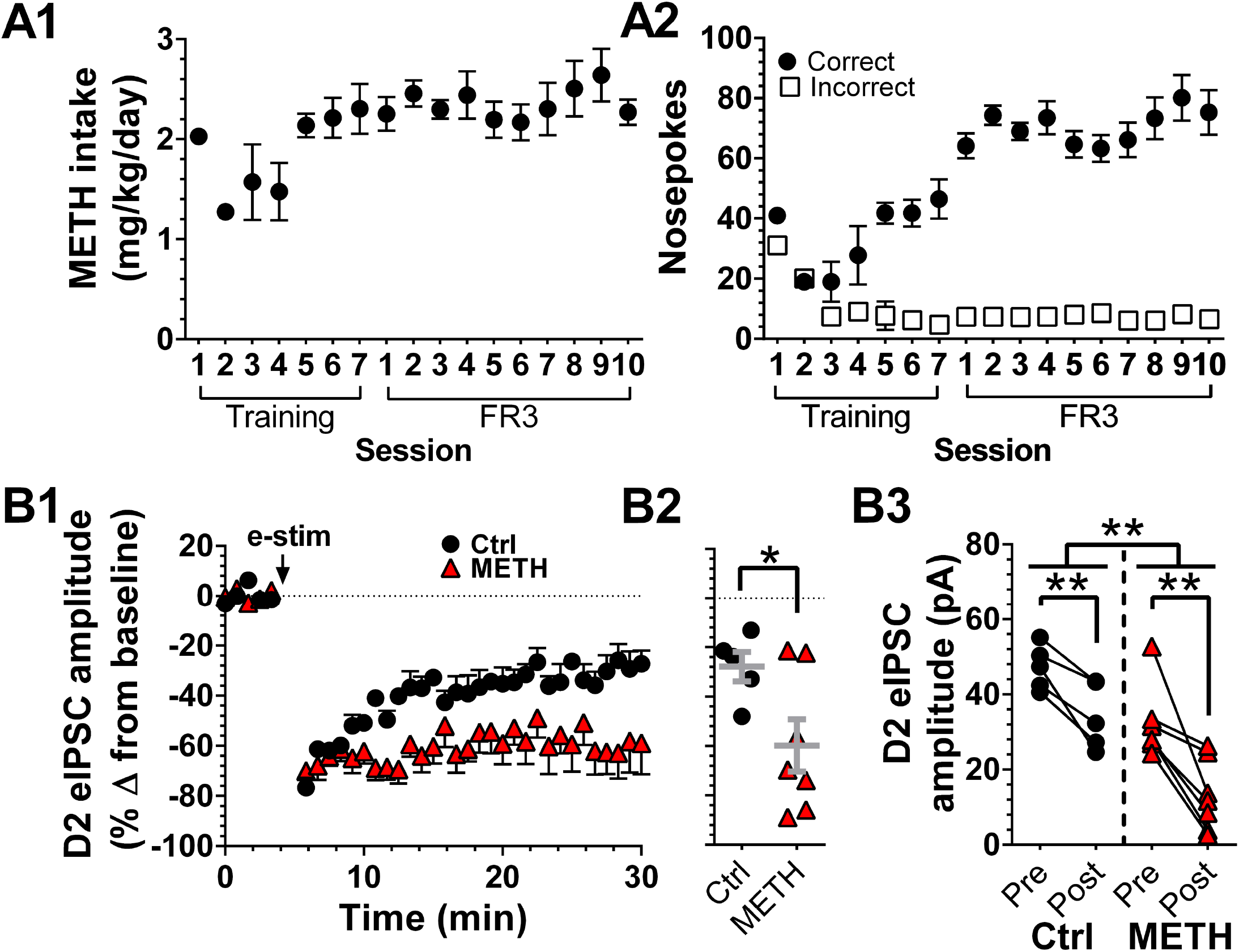
LTD_DA_ is enhanced in mice that previously self-administered methamphetamine. Mice performed methamphetamine (METH) self-administration (i.v.) during daily training sessions (FR1→FR3 schedule), followed by daily maintenance sessions (FR3); average METH intake during each session (A1), and average responding in correct and incorrect nose-poke holes (A2). Effect of electrical stimulation (e-stim, 10 Hz stimulation for 2 min) on D2-eIPSCs recorded from SNc dopamine neurons of mice that self-administered METH (Red triangles) and drug naïve controls (Ctrl, black circles); time-course (B1), average change from baseline (B2), and raw amplitudes (B3). Raw amplitudes are the average of 5 baseline sweeps (Pre) and the 5 sweeps 19-23 minutes following e-stim (Post). *p < 0.05, **p < 0.01

## Discussion

### NT release by dopamine neurons

Our data show that selective optostimulation of dopamine neurons is sufficient to evoke D2 receptor-mediated inhibitory synaptic transmission in these cells, and in the SNc repeated activation of dopamine neurons is sufficient to induce plasticity of that response. This finding is consistent with NT acting as a retrograde transmitter at dendrodendritic synapses between dopamine neurons in the SNc. NT-dependent communication between dopamine neurons meets the criteria of retrograde transmitter release put forth by Regehr and colleagues (2009): NT is present in the postsynaptic cell (Seroogy et al., 1988), disruption of release from the postsynaptic cell prevents signaling, and NT receptors are located presynaptically (Sarret et al., 1998; Tanaka et al., 1990). Further, while local, polysynaptic mechanisms of action are difficult to completely rule out, the lack of effect of antagonizing G_S_-coupled D1 receptors is strong evidence against such a process. Here we show that NT-dependent LTD_DA_ can be inhibited either by 1) blockade of NT receptors, or 2) disruption of NT release through either chelation of calcium or inhibition of V-ATPase. We have previously shown that exposing the presynaptic dopamine neuron to NT by bath application decreases dopamine release (Piccart et al., 2015). Currently available tools do not allow for cell-targeted disruption of NT release from dopamine neurons. NT is not synthesized by dopamine neurons, but enters via sequestration through the high-affinity NT receptor (Castel et al., 1990; Laduron, 1995). Therefore, while somewhat limited by peculiarities of NT physiology, the findings presented here provide strong evidence that NT is released from postsynaptic dopamine neurons and acts as a retrograde messenger to decrease dopamine release presynaptically.

Dopamine neurons also release endocannabinoids which act as retrograde messengers at both GABA and glutamate inputs, or as autocrine signals (Covey et al., 2017). However, a lack of CB-1 endocannabinoid receptors on dopamine neurons suggests that endocannabinoids do not act as retrograde messengers at dendrodendritic dopamine synapses (Julian et al., 2003). Notably, NT signaling via NT type-1 receptors mobilizes endocannabinoid-induced depression of glutamatergic inputs to dopamine neurons (Kortleven et al., 2012). Thus, NT release by dopamine neurons may also induce endocannabinoid release from neighboring dopamine neurons. Taken together, our findings suggest that NT release may act as a dopamine neuron-driven feedback which results in both depression of local dopamine signaling and endocannabinoid-induced plasticity that alters the balance of excitatory and inhibitory inputs to dopamine neurons.

### Dendritic dopamine transmission and methamphetamine self-administration

NT-induced depression of D2 signaling could have a significant impact on reward-related behavior. Burst firing of dopamine neurons, canonically generated by unexpected rewards (Schultz, 1998), strongly drives inhibitory dendrodendritic dopamine signaling and is critical for reward-driven behavior (Beckstead et al., 2004, Zweifel et al., 2009). Dendrodendritic D2 signaling occurs at a timescale that is ideal to terminate burst-firing (Beckstead et al., 2004) and could enhance the specificity of phasic dopamine release. Here we show that D2 current amplitudes in dopamine neurons are reduced by self-administration of the psychostimulant methamphetamine. This could enhance aberrant seeking behavior by circumventing normal inhibitory feedback, blunting the spatial and temporal constraints in dopamine release that typically accompany the presentation of unexpected rewards. Indeed, transgenic mice lacking functional D2Rs show enhanced measures of sensitivity to the rewarding effects of psychostimulants including conditioned place preference, responding on a progressive ratio, acquisition of self-administration, and responding to psychostimulant-paired cues (Bello et al., 2011; de Jong et al., 2015; Holroyd et al., 2015; McCall et al., 2017).

Several forms of synaptic plasticity have previously been described in dopamine neurons following exposure to psychostimulants. Acting through distinct mechanisms, in vivo exposure to cocaine, morphine, and nicotine all induce long-term potentiation of glutamate signaling in dopamine neurons (Brown et al., 2010). Non-contingent cocaine administration causes long-term potentiation of excitatory glutamate signaling (Ungless et al., 2001; Saal et al., 2003; Bellone and Lüscher, 2006) and dysregulation of inhibitory GABA signaling (Liu et al., 2005; Bocklisch et al., 2013), which is thought to increase both the excitability of dopamine neurons and behavioral effects of psychostimulants. The present findings add to this literature by demonstrating that stimulation-induced depression of D2-mediated currents is enhanced by methamphetamine self-administration. This is consistent with previous reports of 2-to-3-fold higher NT levels in the SNc and dorsal striatum following methamphetamine self-administration (Frankel et al., 2011; Hanson et al., 2013). Furthermore, we have shown that blockade or ablation of NT receptors decreases methamphetamine self-administration (Dominguez-Lopez et al., 2018; Dominguez-Lopez et al., 2019). Taken together, this suggests that methamphetamine use enhances NT-induced LTD_DA_, disinhibiting dopamine neurons and driving a further increase in reinforcing effects of the drug.

### NT in the VTA

In the VTA, the endogenous source of NT that drives LTD_DA_ remains unclear. Both electrical stimulation and bath application of NT depress D2-IPSCs in the VTA (Piccart et al., 2015; Stuhrman and Roseberry, 2015). One possibility is that the difference in NT release between SNc and VTA dopamine neurons is due to calcium signaling. NT release in other brain regions is calcium-dependent (Iversen et al., 1978; Kitabgi et al., 1990) and there are substantial differences in calcium binding proteins and ion channel activation in SNc and VTA dopamine neurons (Wolfart et al., 2001, Neuhoff et al., 2002; Rogers, 1992; Liang et al., 1996).

Here we show that NT inputs from both the NAc and the LH to the VTA form synaptic connections with VTA neurons, but generate post synaptic currents in only a small fraction of the neurons recorded and fail to drive LTD_DA_. In agreement with findings reported here of minimal fast-transmitter release from LH and NAc NT-inputs, only 10% of NT expressing terminals form synaptic contacts with dopamine neurons in the VTA (Woulfe and Beaudet, 1992). Furthermore, targeted activation of LH NT-inputs to the VTA increases NT peptide levels but does not increase glutamate or GABA levels when measured by microdialysis (Patterson et al., 2015). To our knowledge this is the first report of NT NAc-inputs forming functional synaptic connections with dopamine neurons. Altogether, it remains to be determined if NT input originating from a single brain region to the VTA drives NT-induced LTD_DA_, and the degree to which upregulation of NT leads to NT release which in turn drives LTD_DA_.

In summary, our findings show that methamphetamine self-administration enhances LTD_DA_ in the SNC, a form of synaptic plasticity that is driven by dopamine neuron activation and subsequent NT release. The long-lasting decrease in dendritic dopamine neurotransmission likely enhances excitability of dopamine neurons and may act as a feed forward mechanism to escalate methamphetamine self-administration behavior.

## Acknowledgements

We would like to thank Dr. Gina Leinninger for providing the *NTS*^*IREScre*^ mice and for comments on the manuscript. We would also like to thank Dr. Stephanie Gantz for comments on the manuscript. This work was supported by National Institutes of Health Grants R01 DA032701 (M.J.B), R01 AG052606 (M.J.B.), the John and Mildred Carlson PhD Scholarship Fund (C.W.T.), and funds from the Presbyterian Health Foundation and the Oklahoma Center for Adult Stem Cell Research, a program of TSET. The authors declare no competing financial interests.

---

**Figure S1.**
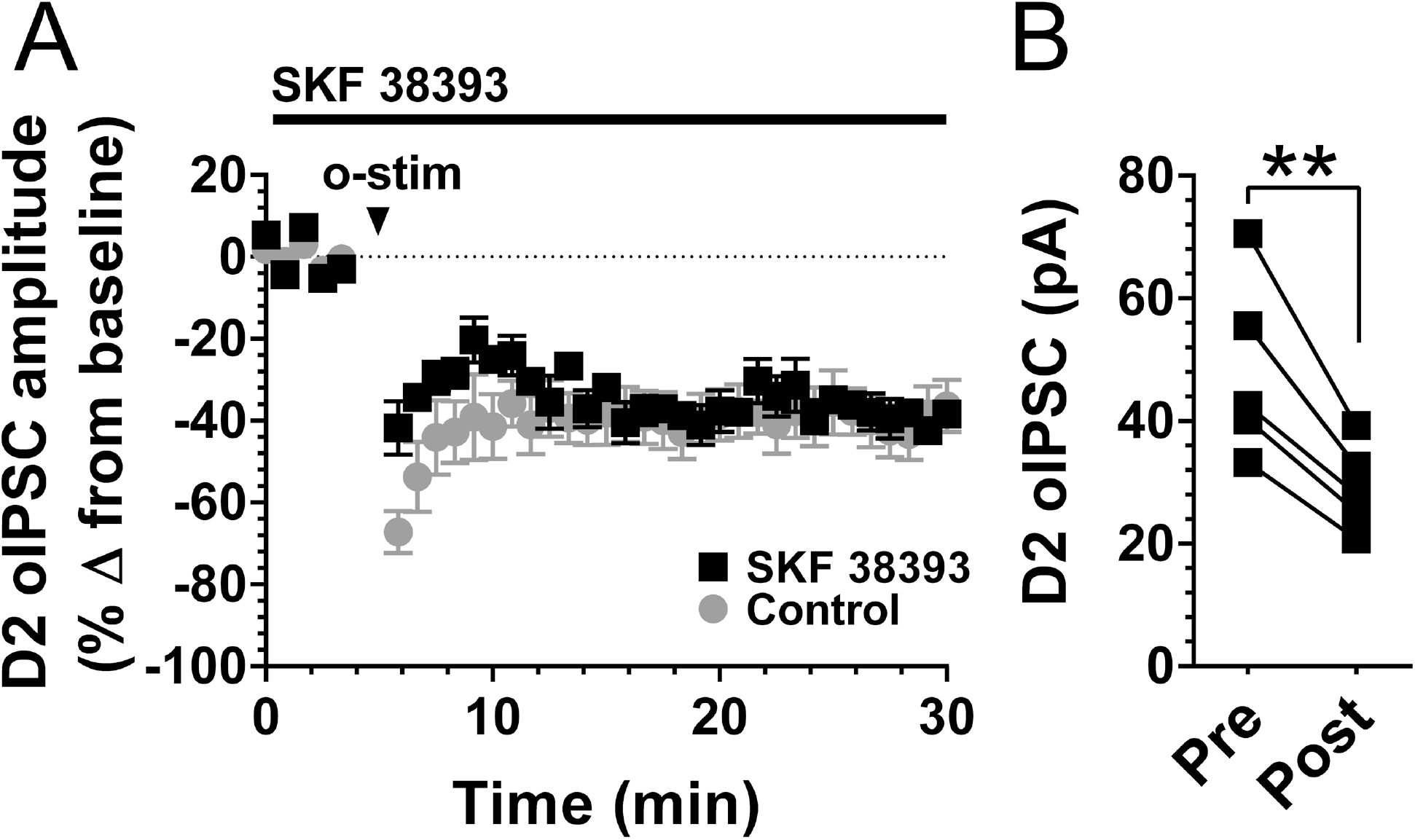
D2-LTD_DA_ is not mediated by dopamine D1 receptors. Effect of dopamine-neuron specific stimulation (o-stim, 10 Hz blue light for 2 min) on D2-oIPSCs recorded from SNc dopamine neurons in aCSF (control, grey circles, data first shown in Figure 1) or in the presence of the dopamine D1 receptor antagonist SKF 38393 (10 µM, SKF, black squares), time-course (A) and raw amplitudes (B). Raw amplitudes are the average of 5 baseline sweeps (Pre) and the 5 sweeps 19-23 minutes following optostimulation (Post).

**Figure S2.**
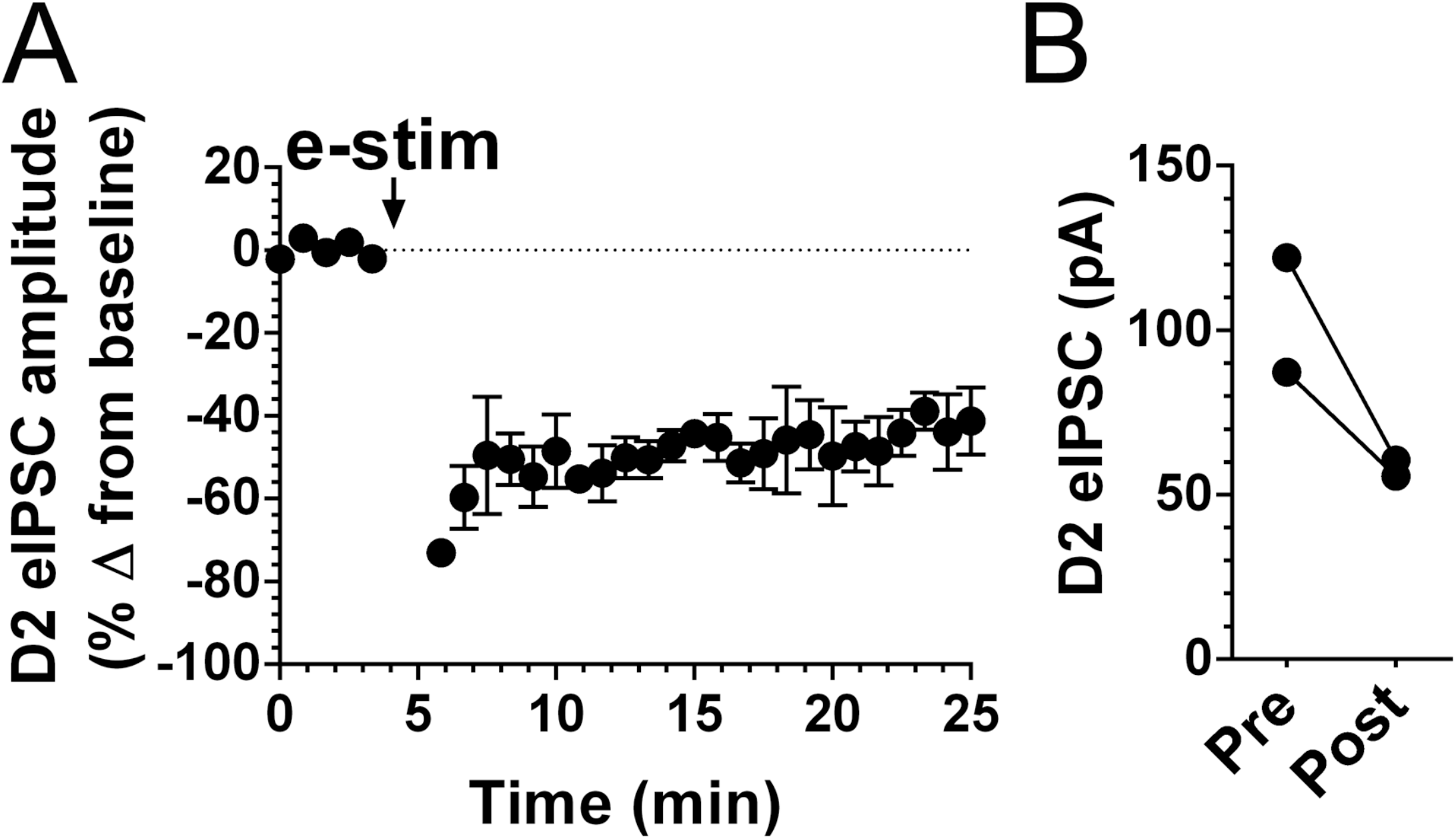
Repeated electrical stimulation drives LTD_DA_ in the VTA. Effect of electrical stimulation (e-stim, 10 Hz stimulation through bipolar electrode for 2 min) on D2-eIPSCs recorded from VTA dopamine neurons (A, B). All raw amplitudes are the average of 5 baseline sweeps (Pre) and the 5 sweeps 19-23 minutes following repeated stimulation (Post).

## Materials and Methods

### Animals

Male and female mice (6 – 52 weeks) were used for all experiments. Genetic lines were obtained from Jackson Labs (C57Bl/6J, DAT-Cre, and Cre-ChR2, Bar Harbor, ME) except for NTS-Cre mice which were generously provided by Dr. Gina Leinninger (Leinninger et al., 2011). Mice were group-housed when littermates were available in polycarbonate chambers with rodent bedding. The vivarium was on a reverse 12/12 light cycle (lights off at 0900). Animal usage was reviewed and approved by Institutional Animal Care and Use Committees at the University of Texas Health Science Center at San Antonio and the Oklahoma Medical Research Foundation.

### Mouse genotyping

DNA was extracted from mouse ear punches using Extract-N-Amp® (Sigma, St. Louis, MO) per the manufacturer’s instructions and used for PCR amplification. Genotyping was performed either on the University of Texas Health Science Center at San Antonio campus using the protocols listed below or performed by Transnetyx (Cordova, TN) using real-time PCR.

DAT-Cre mice (*DAT*^*IREScre*^, *Slc6a3*^tm.1(cre)Bkmn^) and Cre-ChR2 mice (*B6.Cg-Gt(ROSA)26Sor*^*tm32(CAGCOP4*H134R/EYFP)Hze*^/J) were genotyped using a multiplex PCR reaction as per the outlined protocols (https://www2.jax.org/protocolsdb/f?p=116:5:0::NO:5:P5_MASTER_PROTOCOL_ID,P5_JRS_CODE:23107,006660) and (https://www2.jax.org/protocolsdb/f?p=116:5:0::NO:5:P5_MASTER_PROTOCOL_ID,P5_JRS_CODE:28710,024109), respectively. NTS-Cre mice (*NTS*^*IREScre*^, B6;129-*Nts*^*tm1(cre)Mgmj*^/J) were genotyped by multiplex PCR using primers designed by Dr. Richard Palmiter, University of Washington: 5’CAATGGTGCGCCTGCTGGAAG3’, 5’CATGAATGCAAGAAACATCACATCC3’ and 5’TCCAGGAAGATATCCTTGATAACG3’.

### Stereotaxic surgery and vector delivery

Stereotaxic surgeries were conducted using a stereotaxic instrument (Kopf Instruments, Tujunga, CA). Mice were anesthetized with isoflurane kept steady at 1.5-2% using a Somnosuite low-flow anesthesia system (Kent Scientific, Torrington, CT). Access holes were drilled bilaterally in the skull and 33-gauge stainless steel injectors were lowered to one set of the following coordinates in mm: SNc; M/L: 1.25, A/P: −4.7, D/V: −4.7; LH; M/L: 0.9, A/P: −1.1, D/V: −5.25; NAc; M/L: 2.44, A/P: 1.65, D/V: −4.35 @ 20 degrees. At least 3 weeks before electrophysiological experiments, mice were injected with 400 nl of cre-inducible viral vector AAV5-EF1a-DIO-hChR2(H134R)-EYFP-WPRE-pA (University of North Carolina Vector Core, Chapel Hill, NC) synthesized in the lab of Dr. Karl Deisseroth, and mice received injections of 20mg/ml ticarcillin (antibiotic; Sigma-Aldrich) and 0.8 mg/ml ketoprofen (analgesic; Hospira, Lake Forest, IL) post-surgery.

### Catheter implantation surgery

Catheters were constructed in-house using micro-renathane tubing (0.025” o.d., 0.012” i.d; Braintree Scientific, Braintree, MA) with one end attached to a piece of 26-gauge hypodermic tubing (Component Supply, Sparta, TN) bent at a 90-degree angle. The other end of the hypodermic tubing was threaded through the center of a nylon screw. An oval of polyester felt mesh was secured around the base of the screw using silicone medical adhesive (Factor II, Lakeside, AZ). Adhesive was also used to secure hypodermic tubing to the nylon screw with an additional adhesive bead applied approximately 10mm from the unattached end of the micro-renathane tubing to help anchor the catheter during surgery. Aseptic surgeries were performed as described previously (Sharpe et al. 2014). Mice were anesthetized with 2-3% isoflurane using a Somnosuite low-flow anesthesia system. Catheters were placed within a dorsal incision site with the nylon screw perpendicular to the spine. Felt mesh was secured subcutaneously around the dorsal incision while the micro-renathane tubing was threaded subcutaneously to the ventral side. Tubing was inserted into the right jugular vein via ventral incision and anchored using surgical silk sutures (Surgical Specialties, Wyomissing, PA) just below the adhesive bead. Blood was aspirated through the catheter to ensure successful placement. Incision sites were secured with stainless steel wound clips (CellPoint Scientific, Gaithersburg, MD) and the open top of the hypodermic tube was covered by a short piece of heat fused PE20 tubing (Instech Laboratories, Plymouth Meeting, PA) to prevent infection and clotting. Mice received injections of ticarcillin and ketoprofen for pain alleviation and infection prevention post-surgery. Mice were allowed to rest at least 6 days after surgery before beginning daily operant self-administration sessions. Catheters were flushed with heparinized saline (0.2-0.3 ml) (30units/mL; Sagent Pharmaceuticals, Schaumburg, IL) prior to operant sessions.

### Operant self-administration of methamphetamine

Operant sessions were conducted inside modular mouse operant chambers (Lafayette Instruments, Lafayette, IN) with two nose-poke holes on one wall. One hole per chamber was designated as the “correct” nose-poke hole, with nose-pokes resulting in delivery of methamphetamine through surgically implanted jugular catheters. The second nose-poke hole was designated as “incorrect”, and nose-pokes into this hole had no programmed consequences. Daily 2-hour operant sessions were performed as previously described (Dominguez-Lopez et al. 2018). During training mice progressed within-session from a fixed-ratio 1 (FR1) schedule of reinforcement (one nose-poke in the active hole resulting in one infusion of methamphetamine) to a FR3 response requirement. Meeting the response requirement resulted in a 0.1 mg/kg/inf dose of METH (a volume of 12 µl over 2 seconds assuming a 28g adult mouse) directly into their jugular vein. This occurred simultaneously with both a 2 kHz sound cue and illumination of a white-light cue in the chamber’s food receptacle and was followed by a 15 second timeout period during which nose-pokes in either hole had no programmed consequences. Acquisition of self-administration was defined as self-administering at least 8 infusions of METH with at least 70% of responses being made in the correct side nose-poke hole for 2 consecutive sessions. After acquisition, the response requirement to receive one METH infusion was fixed at FR3 and operant sessions continued for a minimum of 9 daily 2-hour sessions followed by 1-3 days in the home cage before *ex vivo* electrophysiological recordings.

### Brain slice electrophysiology

On the day of the experiment, mice were anesthetized with isoflurane and immediately decapitated. Brains were extracted and placed in ice-cold carboxygenated (95% O_2_, 5% CO_2_) artificial cerebral spinal fluid (aCSF) containing the following (in mM): 126 NaCl, 2.5 KCl, 1.2 MgCl_2_, 2.4 CaCl_2_, 1.4 NaH_2_PO_4_, 25 NaHCO_3_, and 11 D-glucose. For Ca^2+^ free experiments CaCl_2_ was replaced with MgCl_2_ (for a total of 3.7 mM) and 0.5 mM EGTA. Kynurenic acid (1 mM) was added to the buffer for the slicing procedure. Horizontal midbrain slices (200 µm) containing the SNc and VTA were obtained using a vibrating microtome (Leica, Wetzlar, Germany). Slices were incubated for at least 30 min at 34-36°C with carboxygenated aCSF that also contained the NMDA receptor antagonist MK-801 (10-20 µM) unless noted otherwise.

Slices were placed in a recording chamber attached to an upright microscope (Nikon Instruments, Tokyo) and maintained at 34-36°C with aCSF perfused at a rate of 1.5 ml/min. Dopamine neurons were visually identified using gradient contrast optics based on their location in relation to the midline, medial lemniscus, and the medial terminal nucleus of the accessory optic tract. Neurons were further identified physiologically by the presence of spontaneous pacemaker firing (1-5 Hz) with wide extracellular waveforms (> 1.1 ms) and a hyperpolarization-activated current (I_H_) of > 100 pA. Recording pipettes (1.5-2.5 MΩ resistance) were constructed from thin wall capillaries (World Precision Instruments) with a PC-10 puller (Narishige International). Whole-cell recordings were obtained using an intracellular solution containing the following (in mM): 115 K-methylsulfate, 20 NaCl, 1.5 MgCl_2_, 10 HEPES, 2 ATP, 0.4 GTP, and either 10 BAPTA or 0.025 EGTA, pH 7.35-7.40, 269-274 mOsm.

D2-IPSCs were evoked during voltage-clamp recordings (holding voltage, −55 mV) using electrical stimulation (D2-eIPSCs, 5 stimulations of 0.5 ms applied at 50 Hz) delivered by bipolar stimulating electrode that was inserted into the slice 100-200 µm caudal to the patched cell in the presence of the following receptor antagonists: picrotoxin (100 µM, GABA_A_), CGP 55845 (100 nM, GABA_B_), DNQX (10 µM, AMPA), hexamethonium (100 µM, nAChR), MPEP hydrochloride (300 nM, mGluR5), and JNJ 16259685 (500 nM, mGluR1). D2-IPSCs were also evoked via blue light (D2-oIPSCs) with 4 stimulations of 5 ms applied at 50 Hz generated either by laser (Optoengine, 100mW max power, 473 nm) and delivered through a fiber optic cable (200 µm core, 0.39 NA, Thorlabs), the tip of which was placed 100-200 µm caudal to the patched cell, or generated by high-power LED (Solis-3C, 20mW max power, Thorlabs) and delivered through the microscope objective after passing through a filter cube (TLV-TE2000, Thorlabs). Both D2-eIPSCs and D2-oIPSCs were evoked once-every 50 seconds. D2-oIPSCs shown in Figure 2 panels D1-2 were recorded in DAT-Cre mice in which ChR2 was expressed by AAV delivered into the SNc (see previous section). For antagonist and inhibition experiments, compounds were either included in the recording pipette and allowed to dialyze for 10 minutes after breaking into the cell, or bath perfused for at least 10 minutes immediately before repeated stimulation and continued throughout the length of the recording.

### Fluorescent labeling and imaging

Brain samples were embedded in OCT cryo-matrix with the coronal fissure oriented to the bottom of the mold. Enough OCT was added to the mold to cover the sample halfway leaving approximately 50% of the sample exposed. The brains were sectioned coronally on a Leica CM3050 cryostat at 40 µm. Each sample was collected at 3 different areas of interest (Franklin and Paxinos, 2008; NAc [1.98 to 0.74mm], LH [-0.58 to −2.54mm], VTA [-2.54 to −3.80mm]). Sections were transferred to a well plate containing a 1X phosphate buffered saline (PBS). Once completed the well plates were sealed with parafilm and stored at 4°C.

Sections were blocked at room temperature for 1 hour with 5% normal donkey serum (Abcam ab7475). Sections were washed in PBS + 0.1% TWEEN-20 then incubated with primary antibodies + 2% normal donkey serum for 22 hours at 4°C. Sections were then washed in PBS and incubated with secondary antibodies for 2 hours at room temperature. After final washes in PBS, sections were mounted on gelatin-coated slides and cover slips were applied using Prolong Diamond anti-fade reagent (Thermo Fisher Scientific). Primary antibodies used were goat anti-GFP (1:500; Novus NB100-1770) for eYFP, and chicken anti-tyrosine-hydroxylase (1:1000; Abcam ab76442). Secondary antibodies used were donkey anti-goat Alexa Fluor 488 (1:750; Jackson ImmunoResearch 705-545-147), and donkey anti-chicken Rhodamine Red-X (1:200; Jackson ImmunoResearch 703-295-155). Primary and secondary antibodies were incubated with PBS + 0.3% triton-x.

Sections were imaged using a Zeiss LSM 710 confocal microscope with a 40x objective. ZEN Microscope Software (Carl Zeiss Microscopy, Thornwood, NY) was used for image acquisition. Final images were obtained at single planes or through maximum intensity flattened z-stacks (1 µm between planes) generated using ImageJ (National Institutes of Health, Bethesda, MD).

### Pharmacological agents

Methamphetamine hydrochloride was a generous gift from the National Institute on Drug Abuse drug supply program (Bethesda, MD). Kynurenic acid, MK-801, DNQX, picrotoxin, dopamine, sulpiride, hexamethonium, SKF38393, concanamycin A, tetrodotoxin, and SR142948 were purchased from Sigma-Aldrich. SR48692, CGP55845, JNJ 16259685, and MPEP hydrochloride were purchased from Tocris Bioscience. The active fragment NT_8-13_ was purchased from American Peptide Company.

### Experimental design and statistical analyses

Data were collected on a Dell computer running Windows 7 using Axograph version 1.5.4 and LabChart (AD Instruments). Statistical analyses were performed using Prism Graphpad version 6. In most cases the effect of repetitive stimulation was analyzed using an average of the 5 sweeps immediately preceding stimulation (sweeps 1-5, pre) and sweeps 31-35 (post). In rare cases where less than 5 sweeps were available, the sweeps within that range (at least 2) were averaged for analysis. Tukey’s post hoc tests were performed subsequent to significant ANOVAs. Data are presented as mean ± SEM. In all cases, α was set a priori at 0.05.

